# Liquid-liquid phase separation within dense-core vesicles in sympathetic adrenal chromaffin cells

**DOI:** 10.1101/2023.11.28.568971

**Authors:** Quanfeng Zhang, Zhaohan Lin, Zhuan Zhou

## Abstract

The neural communication process heavily relies on vesicular release of neurotransmitters and/or neuromodulators primarily from synaptic vesicles (SVs) or dense-core vesicles (DCVs), within neurons and neuroendocrine cells. SVs (∼40 nm) are responsible for releasing classical neurotransmitters, while DCVs (100-500 nm in diameter) release both classic neurotransmitters and neuropeptides. DCVs contain a variety of neuropeptides, hormones, and neurotransmitters, playing crucial roles in diverse physiological processes, such as brain development, neurogenesis, and synaptic plasticity. However, the biogenesis of DCVs and the sorting mechanism of different neuropeptides into DCVs remained largely unknown. Recent studies have revealed that liquid-liquid phase separation (LLPS) of matrix protein-chromogranins plays pivotal roles in the formation of DCVs and the sorting process of neurotransmitters/neuromodulators in endocrine cells. Here we highlight recent advancements in mechanisms of LLPS’s regulation of DCVs ^[1, 2]^, which selectively affect release of some co-transmitters (catecholamines) but not others (i.e. ATP) ^[3]^. We term this phenomenon “1-2-2”: 1 vesicle, 2 transmitters (catecholamine and ATP), 2 release modes (quanta and sub-quanta).

## Introduction

Many components in cells undergo liquid-liquid phase separation (LLPS) to form condensed granules by multivalent interactions through intrinsically disordered regions or binding partners ^[4]^. The LLPS assembly, formed by concentrated molecules that stably exist in a liquid milieu, is a kind of membrane-less compartment (also called biomolecular condensate) and has been found in nucleoli, cytosol, and organelles ^[4-6]^. The LLPS process plays crucial roles in numerous physiological processes, including autophagic degradation, gene expression, enzyme activity modulation, skin barrier formation, vesicle clustering, and postsynaptic densities assembly. However, the complete understanding of granule-liked structures (such as dense-core vesicles (DCVs) in neurons and neuroendocrine cells) remain elusive, and the intriguing question of whether these structures or DCVs are formed through LLPS persists.

In neurons and neuroendocrine cells, regulated secretion occurs primarily from two organelles, synaptic vesicles (SVs) and large DCVs. SVs (∼40 nm in diameter) are responsible for the release of classical neurotransmitters only, while neuropeptides and neurotrophins are transported and secreted *via* large DCVs (100-500 nm in diameter) with a characteristic of electron-dense core. Neuropeptides are essential for the regulation of brain development, neurogenesis, and synaptic plasticity ^[7, 8]^. Abnormal neuropeptide release has been associated with diseases, such as cognitive deficits, psychiatric disorders, drug addiction and obesity ^[8]^. However, the biogenesis of DCVs and the sorting mechanisms of neuropeptides into DCVs remained largely unknown.

DCVs are most abundant in peripheral endocrine cells, such as the sympathetic neuroendocrine adrenal chromaffin cell (ACC, a classical neuroendocrine model for the study of quantal neurotransmitter release ^[9]^) and the insulin-secreting pancreatic β-cells (the release of insulin from pancreatic β-cells plays critical roles in glucose homeostasis, and loss of the first phase of insulin release is considered a hallmark of type 2 diabetes ^[10]^). DCVs also exist in neurons in the central nervous system, where hippocampal and striatal neurons show a large pool of DCVs ranging from 1,400 to 18,000 DCVs per neuron, in correlating with neurite length.

Chromogranins constitute the primary components of the secretory granules or DCVs in the majority of endocrine cells, playing crucial roles in dense-core granulogenesis, sorting of secretory protein cargo, and serving as a reservoir of bioactive peptides ^[11]^. The chromogranin family mainly includes chromogranin A (CgA), chromogranin B (CgB), and secretogranin II (SgII, or CgC), composing of the major matrix proteins in DCVs ^[11]^. The utilization of Protein DisOrder prediction System (PONDR) reveals that all of them display as intrinsic highly disordered proteins, strongly indicating their potential for undergoing LLPS under specific conditions.

Recently, two independent groups reported that chromogranin proteins regulate the biogenesis of DCVs through LLPS in adrenal chromaffin cells ^[1, 2]^. By integrating cutting-edge methods from diverse disciplines such as physiology/electrochemistry and biochemistry/structural biology, we have provided compelling evidence demonstrating that the intra-vesicular matrix protein SgII serves as a pivotal scaffold protein in determining the DCV size *via* LLPS, and substantially regulating quantal release of both catecholamine and neuropeptides in native neuroendocrine ACCs ^[2]^. Moreover, von Blume’s team found that another intravesicular matrix protein CgB underwent LLPS in the lumen of the trans-Golgi network (TGN), and recruited proinsulin to the condensates in a cell-culture model of pancreatic β-cells (INS1 832/13) ^[1]^.

Evidence is mounting that LLPS underlies the formation of membrane-less compartments in cells ^[12]^. The LLPS process concentrates specific components to create a liquid and dynamic membrane-less compartment, facilitating targeted biochemical reactions and playing a crucial role in essential physiological processes ^[5, 12]^. While, LLPS has also been revealed to facilitate the biogenesis of membranous organelles-DCVs ^[1, 2]^. Here we highlight the role of LLPS in recruiting biolipids and the DCVs biogenesis, and tuning neurotransmitters and neuropeptides release.

### LLPS and dense structure under transmission electron microscopy

LLPS in biology was first described in 2002, showing that the hemoglobin could form LLPS droplets *in vitro*, facilitating the nucleation of hemoglobin polymers, while no direct evidence of whether LLPS could be formed *in vivo* at that time ^[13]^. Seven years later, Hyman and his team reported that the P granules undergo LLPS in native cells ^[14]^, thereby establishing LLPS as a physiological phenomenon and solidifying the believe that it occurs naturally within cells. Since then, numerous studies have reported LLPS in various cell types and elucidated their significant roles in related domains, such as modulating reaction kinetics, sequestrating molecules, buffering concentration ^[4, 5]^. In recent years, the combination of the transmission electron microscopy (TEM) and molecular biochemistry has confirmed the dense features observed in pre-synapse (active zone, AZ) and post-synapse (postsynaptic density, PSD) are formed through LLPS of RIM proteins and PSD-related components ^[15]^, respectively. This suggests a potential correlation between the dense-like features observed under TEM and LLPS. The LLPS of synapsin I has been demonstrated to form tight clusters of SVs at synapse, resembling dense structures ^[16]^. Recent studies have also shown that DCV matrix proteins CgA, CgB and SgII undergo LLPS, which contributes to vesicular cargo package and vesicle volume determination ^[1, 2]^. These collective findings strongly support the significance of LLPS in biological processes and its association with dense-like features observed under TEM.

### Chromogranins-LLPS and lipids

*In vivo* and *in vitro* studies have demonstrated that protein LLPS assembly can recruit multiple proteins or nucleic acids (DNA/RNA) into the the LLPS bodies, forming membrane-free biomolecular dynamic co-condensates that execute complex and diverse biological functions. Examples include PSD-95/Shank3/Homer3/GKAP in the post-synapse, RIM/RIM-BP in the pre-synapse, and DNA-protein co-condensation in chromatin formation and transcription initiation ^[15]^. However, little is known about the relationship between LLPS and lipids. Pietro De Camilli’s team demonstrated in 2018 that synapsin I undergoes LLPS and recruits SVs locally, leading to the formation of SV clusters at presynaptic sites ^[16]^. In this context, the lipid components of SVs can form clusters through the synapsin I-mediated LLPS. The recruitment mechanism between LLPS and SVs is facilitated by the synaptic-vesicle binding region (residues: 113-420) of synapsin I. This region acts as an anchor on the SV membrane, enabling the incorporation of SVs into LLPS.

Recently, through the integration of electrochemical carbon fiber electrodes (CFE) recording (detection of single DCV catecholamine release), biochemistry (mammalian protein purification), structural biology (cryo-electron microscopic analysis of helical reconstruction), and imaging (correlative light-electron microscopy, TEM and confocal imaging), our group has discovered that SgII, a DCV matrix protein lacking a transmembrane domain, can undergo LLPS both *in vitro* and *in vivo*. Furthermore, we have observed that its LLPS assembly is capable of recruiting bio-lipids to form lipid-coated SgII droplets, which plays a vital role in regulating the biogenesis of DCVs and further modulates the release of quantal neurotransmitters and neuropeptides from native neuroendocrine cells ^[2]^.

First, we observed that SgII exhibits pH-dependent polymerization (monomers at pH 5.5; dimers/polymers at pH 7.5). The helical structure of polymerized SgII was analyzed using cryo-electron microscopy (cryo-EM) (**Fig. 1A**, see also ^[2]^). Further, under appropriate pH conditions mimicking the native environment of the trans-Golgi network (TGN) lumen, SgII undergoes LLPS, leading to the formation of droplets both *in vitro* and *in vivo* (**Fig. 1B**, see also ^[2]^). These SgII-LLPS droplets can recruit reconstituted bio-lipids, facilitating the biogenesis of DCVs (**Fig. 1B**, see also ^[2]^). In addition, by employing correlative light-electron microscopy (CLEM) and electrochemical CFE recording techniques to detect single DCVs release from native ACCs, we revealed the pivotal physiological role of SgII-LLPS: knockdown of SgII significantly reduced the quantal neurotransmitter release by affecting DCV size, which can be rescued differently by various truncations of SgII with distinct degrees of LLPS capability (**Fig. 1C**, see also ^[2]^). Secretogranin III (SgIII), another member in chromogranin family, has been proved to dissolve SgII-mediated LLPS *in vitro* and reduce the volume of DCVs in native ACCs through overexpression (**Fig. 1D**) ^[2]^. Moreover, using CFE recording, we observed that the reduction in DCV volume caused by SgIII overexpression leads to a reduction in ACC quantal size ^[2]^. These findings highlight the unique role of SgII as an intravesicular matrix protein involved in LLPS and provide novel insights into how SgII regulates DCV biogenesis and the quantal release of neurotransmitters and neuropeptides (**Fig. 1D**, see also ^[2]^).

**Fig. 1.**
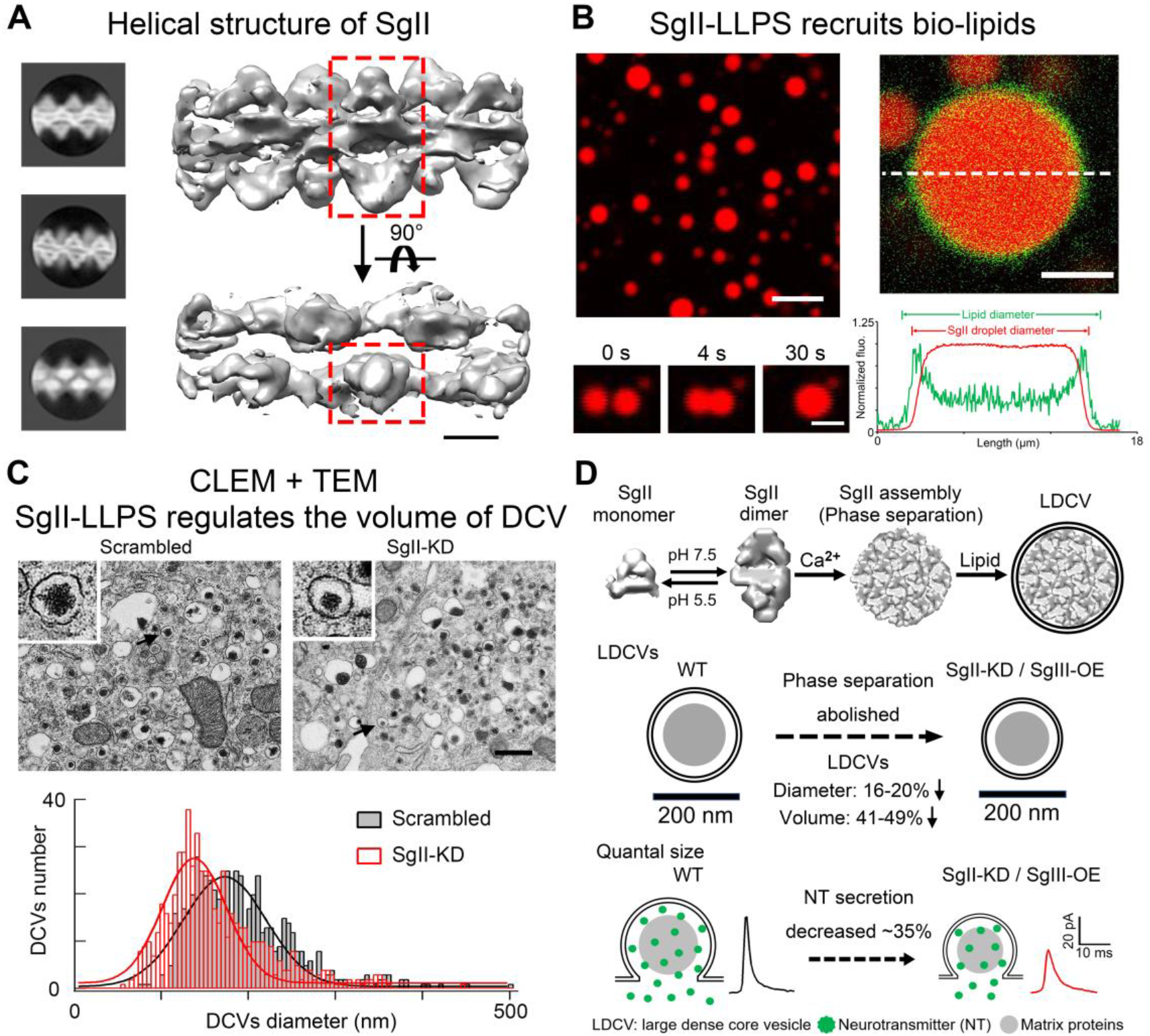
SgII regulates the DCV size and quantal neurotransmitter release *via* LLPS.^[2]^. A. Representative classes from 2D-classification of SgII filament structure under cryo-EM (left). Reconstructed 3D model of SgII helical structure. The one dimeric subunit is outlined by red dashed lines (scale bar, 50 Å) (right). B. SgII forms droplet structures with Ca^2+^ under confocal microscopy (scale bar, 10 μm) (upper left). The SgII-mediated droplets can fuse with each other to form a bigger droplet, which exhibits LLPS behavior (scale bar, 3 μm) (lower left). The SgII-mediated LLPS droplet can be coated by bio-lipids (upper right). Plot profiles of normalized fluorescent intensity between SgII-LLPS and bio-lipid under white line (scale bar, 5 μm) (lower right). C. Representative transmission EM micrographs of DCVs from scrambled and SgII-KD ACCs (scale bar, 500 nm) (upper). The distribution of DCV diameters shows smaller DCVs in SgII-KD ACCs than scrambled ACCs (lower). D. The mechanism of SgII-LLPS determining DCV volume and quantal release in mammalian neuroendocrine cells.

To understand the role of SgII-mediated LLPS in regulating the volume of DCVs, the relationship between LLPS assembly and DCV membrane need to be clarified. Based on current knowledge, matrix proteins are packaged within the TGN prior to DCV budding, suggesting that the TGN serves as an optimal site for SgII-mediated LLPS influencing DCV volume. I*n vitro* experiments have demonstrated that reconstructed bio-lipids (comprising 250 μM DSPC, 150 μM DOPE, 50 μM SoyPI, 5 μM DMB PEG and 5 μM NBD PC) can effectively coat SgII-mediated LLPS assemblies, mimicing the physiological composition of TGN membrane (**Fig. 2A**) ^[2]^. While circular coated bio-lipids features have not been observed in groups of Ca^2+^ with bio-lipids (**Fig. 2B**), SgII alone with bio-lipids (**Fig. 2C**) and SgII-LLPS assemblies with 25% ethanol (bio-lipids buffer) (**Fig. 2D**). Without a membrane-anchoring domain, the SgII-mediated LLPS can also recruit lipids as well, indicating a distinct recruitment mechanism compared to synapsin I. It is possible that lipids can directly interact with SgII. Based on hydrophobicity analysis, most parts of SgII are hydrophilic, suggesting that cations (Ca^2+^ and Mg^2+^) serve as the driving force for LLPS ^[2]^. However, some hydrophobic regions may still interact with lipids. Considering physiological pH levels and appropriate concentrations of SgII and Ca^2+^, it is believed that SgII undergoes LLPS in TGN ^[2]^ and forms LLPS assembly (**Fig. 2E**). Once these assemblies come into contact with the TGN membrane, they may recruit membrane components and participate in the DCV biogenesis while regulating its volume (**Fig. 2E**). This protein-lipid interaction involving LLPS presents a novel pattern of LLPS-mediated lipid recruitment that may be involved in several cases related to membrane-containing organelles or the plasma membrane.

**Fig. 2.**
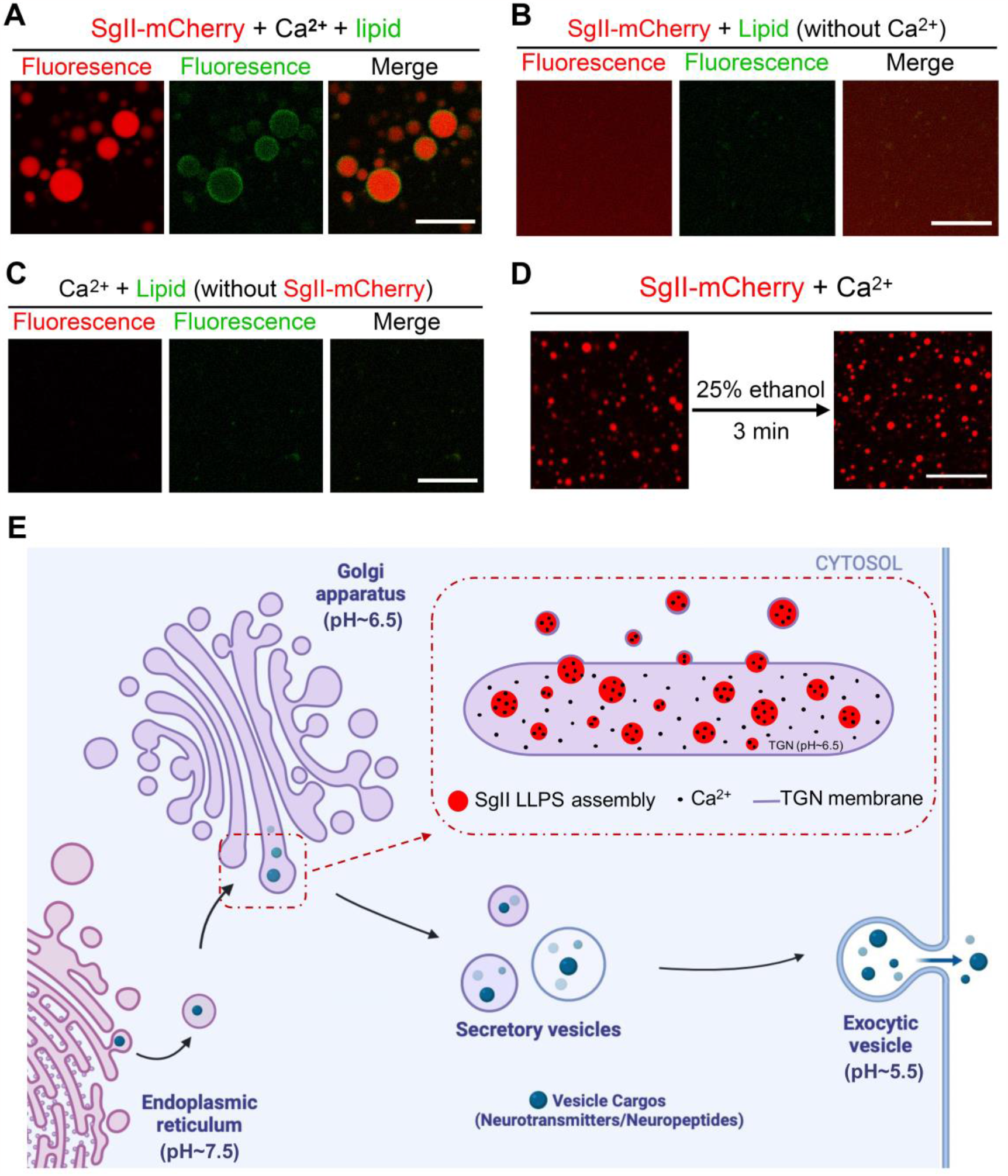
The SgII LLPS assembly recruits bio-lipids, mimicking the biogenesis of DCVs. ^[2]^. A. SgII LLPS droplets can be coated by reconstituted bio-lipids under confocal imaging (scale bar, 5 μm). B. SgII LLPS droplets are not observed with lipid alone (without Ca^2+^) (scale bar, 20 μm). C. Circular lipid structures are not observed with Ca^2+^ alone (without SgII) (scale bar, 20 μm). D. SgII LLPS droplets are not influenced by 25% (v/v) ethanol (bio-lipids buffer) (scale bar, 20 μm). E. Proposed model of how SgII-LLPS regulates DCV volume during vesicle biogenesis. We propose that SgII undergoes LLPS in TGN with proper situations (sufficient concentrations of SgII and Ca^2+^, suitable pH environment). Next, the LLPS assemblies participate in DCV initial formation through touching and binding the TGN membrane, subsequently budding off to form the immature vesicle. Finally, during the lumen acidification from immature (pH 6.5) to mature vesicle (pH 5.5), the inside SgII-mediated LLPS will disperse.

### Neurotransmitters release and chromogranins-LLPS

The amount of neurotransmitter release from individual vesicles triggered by Ca^2+^ influx is defined as quantal size, which determines the strength and speed of synaptic transmission. The micro electrochemical CFE is a real-time and highly sensitive tool for real-time monitoring of single DCV catecholamine quantal release through electro-oxidation in ACCs ^[3, 17]^. It has been established that proteins/ions outside of vesicles can regulate the amperometric quantal size of DCVs by affecting vesicle fusion pore dilation, including the G protein-coupled receptor βγ, the SNARE complex, dynamin, and Ca^2+^ [^3, 18-20]^. Additionally, changes in the size/volume of DCVs are associated with corresponding change of quantal size. Recently, our research revealed that CgA is an essential factor for the formation of sub-quantal release by binding to catecholamines (but not ATP) and limiting their diffusion flux from a small vesicle fusion pore, thus producing the phenomena “1-2-2”: 1 vesicle, 2 transmitters (catecholamine and ATP), 2 release modes (quanta and sub-quanta) ^[3]^, suggesting the critical roles of intra-vesicular matrix proteins such as chromogranins (including SgII) on DCV quantal secretion ^[2]^. Furthermore, we have hypothesized that chromogranins may undergo LLPS to regulate the release of neurotransmitters in DCVs ^[3]^. By employing biochemical and structural biology techniques, our recent study has proved that the LLPS phenomenon of SgII (one of the matrix protein) and demonstrated how a small amount of the matrix protein SgII can dictate the size of DCVs. Our findings together demonstrated that SgII serves as a scaffold protein for DCV in ACCs to determine DCV size and impact the quantal size (**Fig. 1D**). The recruitment of bio-lipids mediated by chromogranins-LLPS promotes the formation of vesicles which offers novel perspectives on LLPS.

### Final Remarks

DCVs play a crucial role in the sustained (sub-quantal) release of neuropeptides and hormone in many endocrine and neuroendocrine cells, as well as neurons, to regulate numerous essential physiological processes ^[8]^. Compared to SVs (∼40 nm), DCVs (100-500 nm in diameter, with an electron dense-core and a larger vesicle volume under TEM) exhibit stronger enrichment and storage capabilities for neurotransmitters and neuropeptides, primarily due to the presence of intracellular matrix proteins known as chromogranins including CgA and SgII within DCVs. CgA binding of catecholamines helps maintain the high concentration of catecholamines (0.8-1 M) within DCVs of ACC, facilitating sub-quantal release of catecholamines through pH-dependent matrix dissociation during vesicle fusion opening process ^[3]^. Additionally, SgII acts as a scaffold protein for ACC’s DCVs by determining vesicle size through LLPS recruitment of bio-lipids ^[2]^. It further increases the amount of neurotransmitter loaded into the vesicle, thereby affecting quantal release of catecholamines and neuropeptides from individual DCVs ^[2]^. The recent works from two groups have proved the causal relationship between matrix protein SgII and CgB for LLPS and their regulation on DCV biogenesis, sorting and transmission of neurotransmitters (catecholamines) & neuropeptides (insulins) ^[1, 2]^. Future work should address the role of LLPS in mediating differential co-release of two native neurotransmitters from a single DCV (“1-2-2” phenomena: 1 vesicle, 2 transmitters (catecholamine and ATP), 2 release modes (quanta and sub-quanta)).

## Author Contributions

Q.F.Z, Z.H.L and Z. Z wrote the manuscript.

## Acknowledgments

This work was supported by National Natural Science Foundation of China (31930061, 21790394, 82000757, 31761133016, 31821091, 31330024, 31171026, 31327901, 32171233, 31670843 and 21790390), the National Key Research and Development Program of China (2016YFA0500401), the National Postdoctoral Program for Innovative Talents (BX20190012), and the China Postdoctoral Science Foundation (2020M670029).

## References

[1] Parchure A, Tian M, Stalder D, Boyer CK, Bearrows SC, Rohli KE, et al. Liquid-liquid phase separation facilitates the biogenesis of secretory storage granules. J Cell Biol 2022, 221.

[2] Lin Z, Li Y, Hang Y, Wang C, Liu B, Li J, et al. Tuning the Size of Large Dense-Core Vesicles and Quantal Neurotransmitter Release via Secretogranin II Liquid–Liquid Phase Separation. Advanced Science 2022: 2202263.

[3] Zhang Q, Liu B, Wu Q, Liu B, Li Y, Sun S, et al. Differential Co-release of Two Neurotransmitters from a Vesicle Fusion Pore in Mammalian Adrenal Chromaffin Cells. Neuron 2019, 102: 173–183 e174.

[4] Banani SF, Lee HO, Hyman AA, Rosen MK. Biomolecular condensates: organizers of cellular biochemistry. Nat Rev Mol Cell Biol 2017, 18: 285–298.

[5] Shin Y, Brangwynne CP. Liquid phase condensation in cell physiology and disease. Science 2017, 357.

[6] Ryan VH, Fawzi NL. Physiological, Pathological, and Targetable Membraneless Organelles in Neurons. Trends Neurosci 2019, 42: 693–708.

[7] Park H, Poo MM. Neurotrophin regulation of neural circuit development and function. Nat Rev Neurosci 2013, 14: 7–23.

[8] van den Pol AN. Neuropeptide transmission in brain circuits. Neuron 2012, 76: 98–115.

[9] Chow RH, von Ruden L, Neher E. Delay in vesicle fusion revealed by electrochemical monitoring of single secretory events in adrenal chromaffin cells. Nature 1992, 356: 60–63.

[10] Rorsman P, Renstrom E. Insulin granule dynamics in pancreatic beta cells. Diabetologia 2003, 46: 1029–1045.

[11] Taupenot L, Harper KL, O’Connor DT. The chromogranin-secretogranin family. N Engl J Med 2003, 348: 1134–1149.

[12] Alberti S, Gladfelter A, Mittag T. Considerations and Challenges in Studying Liquid-Liquid Phase Separation and Biomolecular Condensates. Cell 2019, 176: 419–434.

[13] Galkin O, Chen K, Nagel RL, Hirsch RE, Vekilov PG. Liquid-liquid separation in solutions of normal and sickle cell hemoglobin. Proc Natl Acad Sci U S A 2002, 99: 8479–8483.

[14] Brangwynne CP, Eckmann CR, Courson DS, Rybarska A, Hoege C, Gharakhani J, et al. Germline P granules are liquid droplets that localize by controlled dissolution/condensation. Science 2009, 324: 1729–1732.

[15] Chen X, Wu X, Wu H, Zhang M. Phase separation at the synapse. Nat Neurosci 2020, 23: 301–310.

[16] Milovanovic D, Wu Y, Bian X, De Camilli P. A liquid phase of synapsin and lipid vesicles. Science 2018, 361: 604–607.

[17] Wightman RM, Jankowski JA, Kennedy RT, Kawagoe KT, Schroeder TJ, Leszczyszyn DJ, et al. Temporally resolved catecholamine spikes correspond to single vesicle release from individual chromaffin cells. Proc Natl Acad Sci U S A 1991, 88: 10754–10758.

[18] Blackmer T, Larsen EC, Bartleson C, Kowalchyk JA, Yoon EJ, Preininger AM, et al. G protein betagamma directly regulates SNARE protein fusion machinery for secretory granule exocytosis. Nat Neurosci 2005, 8: 421–425.

[19] Chen XK, Wang LC, Zhou Y, Cai Q, Prakriya M, Duan KL, et al. Activation of GPCRs modulates quantal size in chromaffin cells through G(betagamma) and PKC. Nat Neurosci 2005, 8: 1160–1168.

[20] Shin W, Ge L, Arpino G, Villarreal SA, Hamid E, Liu H, et al. Visualization of Membrane Pore in Live Cells Reveals a Dynamic-Pore Theory Governing Fusion and Endocytosis. Cell 2018, 173: 934–945 e912.

